# Computational design of artificial supply networks for engineered human tissue

**DOI:** 10.1101/2025.10.21.683642

**Authors:** Henning Bonart, Pramodt Srinivasula, Ulrike A. Nuber, Steffen Hardt

## Abstract

The development of large-scale, three-dimensional human tissues is crucial for various applications in therapeutic tissue engineering, disease modeling, and drug testing. However, due to the diffusion limit of oxygen, the lack of functional vascular networks is a significant limitation in maintaining these engineered tissues in the laboratory. To address this challenge, we present a systematic, model-based design process for artificial supply networks that can ensure a sufficient supply of oxygen and nutrients to engineered human tissue. Our approach combines mathematical models of fluid dynamics, cell metabolism, and network properties to identify key parameters influencing the supply performance. We demonstrate the applicability and possibilities of this design process by simulating different network structures, including cuboid and rhombic dodecahedral honeycombs, under various conditions. Our results show that the structure of the artificial supply network, oxygen concentration, and solute flow within the network strongly influence cellular metabolic activity and viability. We also examine the effects of non-uniform cell density, channel blockage, and long channel length on the oxygen distribution inside the cell-containing tissue compartment. Our findings highlight the importance of considering these factors in the design of artificial supply networks for large-scale engineered human tissues. This study provides a promising approach for quickly exploring the vast design space of possible network structures under different conditions for desired cell and tissue states, ultimately contributing to the development of more efficient and effective tissue engineering strategies.

## 1. Introduction

Human tissue is produced and maintained in laboratories for research on diseases, for toxicity and efficacy testing of substances, and for therapeutic tissue engineering [1, 2]. To imitate key functional and structural characteristics of natural human tissue as closely as possible, these applications require human tissue of sufficient size. For certain applications, including tissue replacement therapies, two-dimensional cell cultures and single small three-dimensional cell culture models like spheroids or more complex stem cell-based organoids with diameters of a few hundred micrometers do not meet critical size requirements [3– 6]. Already the smallest functional units of human tissues have dimensions in the mm to cm range and a three-dimensional (3D) structure. For example, colonic crypts have a nearly cylindrical shape with a submillimeter size [7]. Nephrons, the functional units of a kidney, possess submillimeter-sized glomeruli (head of the geometry) attached to tubuli of multiple centimeters in length (tail of the geometry) [8]. Other examples of functional human tissue units are liver lobules which are about a millimeter in diameter as well as height (see references in [9]). Similarly, neurons of brain tissue can have axons, specialized cellular processes, of submillimeter to multiple centimeters in length, many of them with an elaborate 3D structure.

The diffusive transport of oxygen through human tissue combined with its metabolic consumption by cells results in a spatial nonuniformity of the oxygen concentration and a certain length scale over which oxygen is consumed in the tissue. Hence, when larger tissue constructs are placed in a liquid culture medium to supply oxygen by diffusion from this external medium, dead or dying cells are found at the tissue center in case of tissue diameters greater than a few hundred micrometers [10–14]. Nature’s design to overcome this limitation is the hierarchical vascular network, which also contains mesh-like areas, for example capillary beds [15–17]. *In vivo*, the composition of blood, the morphology of the vascular tree, and capillary types of different permeability are such that oxygen and other species are provided and removed according to the metabolic requirements of a tissue. The distance between capillaries, the smallest blood vessel types, which mediate species exchange to and from the tissue, is only a few ten to hundred micrometers [18–21]. This distance is adapted to the cell density and the extent of cellular metabolic activity. Furthermore, static culturing methods, such as suspension cultures of spheroids or scaffold/hydrogel-embedded 3D cell cultures, cannot reproduce the dynamic nutrient transport of *in vivo* tissues [22, 23].

The lack of vascular systems that transport oxygen, nutrients, and metabolic waste products to or from cells is the major limitation that prevents constructing large 3D human tissues with diameters of more than several hundred µm (see reviews [24–26] and references therein). Therefore, various approaches to providing laboratory-generated human tissues with a natural vasculature or with an artificial supply network, fulfilling species transport and exchange functions similar to a natural vascular system, have been researched. Building a natural vasculature in human 3D tissue such as organoids through co-culturing endothelial cells, which constitute the inner lining of blood vessels, together with other vasculature-forming cells, is an ongoing pursuit in tissue engineering [27–29]. An alternative is to build supply networks from synthetic materials to serve as a vasculature for human tissue. This can for example be achieved by pre-fabrication, e.g., via additive manufacturing, or by sacrificial materials, which leave hollow channels after their removal [16, 30–32].

Compared to exclusively naturally grown vascular networks, artificial supply networks provide several major advantages. Firstly, it takes a long time for blood vessels to grow into a tissue. An average growth rate of only ∼ 5 *µ*m per hour has been reported for newly developing microvessels [33] Secondly, although endothelial cells can form microvascular networks when combined with other cell types within 3D cell cultures, larger blood vessel types are not generated in this way. By contrast, an artificial supply network can - irrespective of its channel diameters - be integrated during the tissue building process so that the incorporated cells are never insufficiently supplied for prolonged periods. Importantly, artificial supply networks can be combined with natural blood vessels to supply a tissue created in the laboratory, using anastomoses that are established between endothelialized channels of supply networks and natural capillaries formed in human 3D tissue [34–37] In addition, supply networks can also be integrated in a tissue *in vivo* through surgical anastomosis with blood vessels of medium to large size [34, 38]. Furthermore, artificial supply networks offer the integration of sensor and actuator elements in and on network walls for the targeted monitoring and control of cells in engineered human tissue. Finally, artificial supply networks have a wider range of applicability compared to the natural ones and they can be systematically designed to fulfill specific goals under certain constraints. Importantly, evident differences between the species transport through supply networks for engineered human tissue and *in vivo* vascular networks need to be considered when designing artificial supply networks. Blood, a non-Newtonian fluid, is circulated through deformable natural vessels and the dissolved oxygen concentration in blood is about more than 30 times higher than that in an aqueous cell culture medium, due to oxygen binding to hemoglobin in red blood cells.

The structure of the supply network and the supply of metabolic species from it strongly influence cellular activity outside the network. The cellular metabolic consumption of oxygen along its diffusive path in the tissue results in a spatial nonuniformity of the oxygen concentration. This leads to variability in cell proliferation, viability, and phenotype. Hence, to ensure a metabolic similarity of engineered human tissue to *in vivo* conditions, it is essential to systematically study the transport of metabolic species through the artificial supply networks and the cell-containing tissue compartment in combination with the metabolism of the cells. Here, mathematical and computational methods can contribute decisively to understand and improve such complex engineered human tissue [39]. In [40] Kirchhoff’s laws where utilized to compute the flow distribution through a three-dimensional capillary mesh network coupled to tissue diffusion equations including Michaelis-Menten kinetics, while important design aspects like channel wall permeability, inlet oxygen concentration, or the tissue-to-network volume ratio were not considered. Recently, a combination of constrained constructive optimization, multifidelity CFD, and direct 3D bio-fabrication was demonstrated for the design of hierarchic branching networks [41]. However, this study evaluated tissue perfusion only through approximate analytical solute depletion models and did not couple network flow dynamics to a spatially resolved cellular metabolism model, providing no direct design guidance on wall permeability, inlet oxygenation, or cell density relative to a hypoxic threshold. Besides, several further studies have applied computational oxygen transport models to channel network problems, including [42] to optimize the microchannel geometry in a vascular graft wall, [43] to compare bifurcating tree designs, [44] to design a biomimetic liver scaffold, and [45] to derive bioreactor design criteria. However, a systematic, network-level, multi-parameter design of supply networks by coupling flow and transport inside the channels, wall permeability, as well as metabolic reactions and diffusion in tissue compartments is still missing.

Regarding the use of artificial supply networks for 3D tissues generated in the laboratory, although there has been significant progress concerning materials and fabrication techniques, as well as model-based design, critical design aspects to ensure a sufficient oxygen and nutrient supply as well as waste removal in engineered human tissue as a whole remain underexplored. Therefore, we present a systematic design process based on a coupled model including cell metabolism and transport, both in the cell-containing tissue compartment and in the network. Based on this integrative model we identify key parameters influencing the design of artificial supply networks, and demonstrate examples of relevant design outcomes.

## 2. Systematic design process

### 2.1. General approach

The design of artificial supply networks for engineered human tissue is an interdisciplinary and iterative effort involving a coordinated interplay between biology, fluid dynamics, and material science. The possible designs are mainly influenced by the density and metabolism of the cell types, the properties of the cell environment, for example hydrogels used as extracellular material, the flow and transport inside the supply networks, and the type of network fabrication. Naturally, this results in a vast parameter space, making a purely experimental approach based on prototyping inefficient. In contrast, a systematic, computer-assisted design process with mathematical models based on fluid dynamics, tissue metabolism and network properties can identify key parameters influencing the supply performance. Further, such a design process reduces the number of prototypes and experiments needed to identify a suitable or even optimal solution.

In Figure 1 we present a systematic design process for artificial supply networks. To start the process, we assume that the choice of cell types with defined metabolic profile and desired tissue status, e.g., normoxic conditions above a defined hypoxic boundary, chemical kinetics, and cell densities, has already been fixed (yellow box in Figure 1)). In the next step, several design parameters have to be set in a way that the desired cell/tissue status is maintained (green boxes). Here we apply computer experiments based on models of a) the cell environment (e.g., hydrogel that contributes to net oxygen diffusivity), b) the channel properties (e.g., length or mass transfer area), and c) the flow conditions, e.g., volume flow rate or oxygen and nutrient concentrations (dashed box, large). Note that the cell environment can vary over space and time to enable the cells to proliferate and self-organize into a specific larger 3D tissue construct. The evaluation of the model for every point in parameter space is fast and cheap, which allows to iteratively try out different combinations and formulate a promising design proposal of an artificial supply network. Subsequently, a physical prototype of the design is manufactured and the cell/tissue status is experimentally verified (blue box). If the results match the desired cell/tissue status, the design process terminates successfully. However, in case of a mismatch, the model parameters or even the model itself have to be adjusted to better represent the complex reality, and the model-based design loop is repeated.

**Figure 1:**
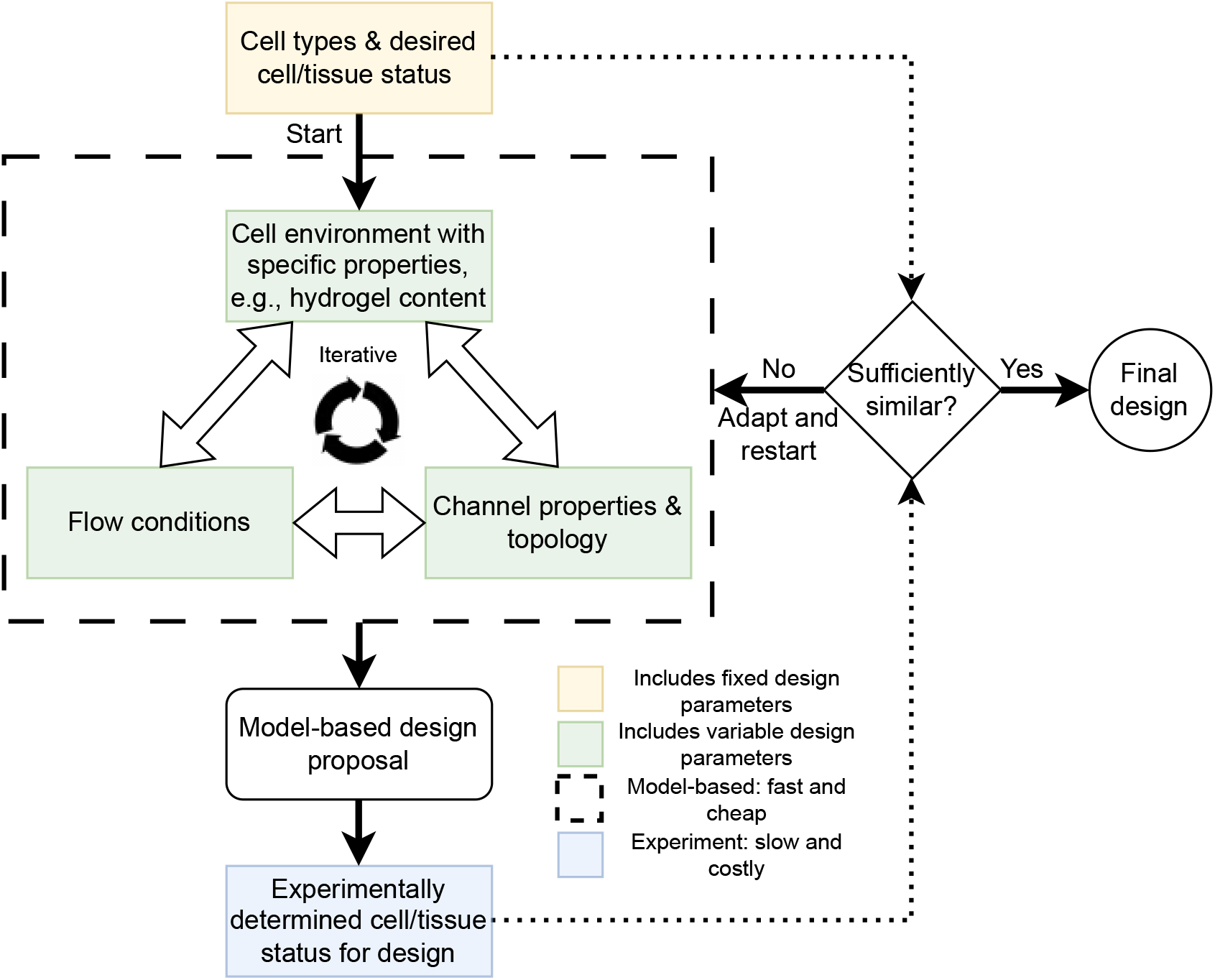
Systematic, computer-assisted design process for artificial supply networks.

### 2.2. Modelling tissue and supply network

We consider the metabolic activity of human cells inside cell-containing tissue compartments as well as the transport between these compartments and the artificial supply network as a coupled system, see Figure 2. The simulation domain is decomposed into the channel interior given by Ω_*c*_, the wall of the channels given by Ω_*w*_, and cell-containing tissue compartment given by Ω_*t*_.

**Figure 2:**
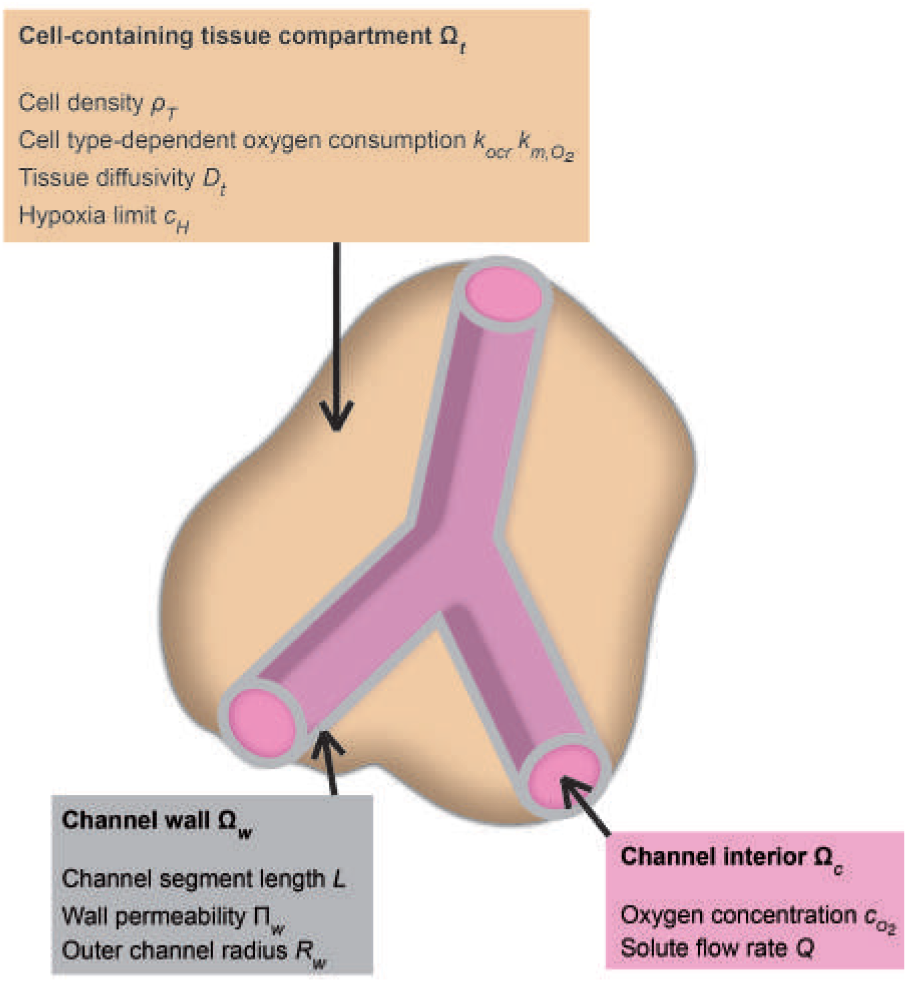
Three coupled regions (channel interior Ω_*c*_, channel wall Ω_*w*_, and cell-containing tissue compartment Ω_*t*_) in the model and the most important parameters describing cell metabolism (cell density *ρ*_*T*_, cell type-dependent oxygen consumption *k*_*ocr*_ and 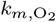, hypoxia limit *c*_*H*_), species transport (tissue diffusivity *D*_*t*_, oxygen concentration inside the channel 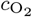, solute flow rate *Q*), and channel properties (channel segment length *L*, wall permeability Π_*w*_, outer channel radius *R*_*w*_).

We focus on aerobic metabolism, since it is most common among normal tissues, and sufficient data for comparison with past and future studies exist. We assume that an aqueous cell culture medium with dissolved oxygen is pumped through the network channels. Apart from saturating the medium with oxygen from air at atmospheric pressure, we also consider conditions up to the maximal oxygen solubility in the aqueous medium at 37 ^◦^C. This can for example be achieved by using pure oxygen at atmospheric pressure or by increased pressures for air. Additionally, a sufficient contact time and surface area between the gas and liquid phase needs to be achieved to reach saturation, e.g., through a membrane saturator [46]. The cell environment, which contributes to the oxygen diffusion inside the cell-containing tissue compartment, can be a natural extracellular matrix, produced by the cells themselves, or a synthetic matrix, e.g., a hydrogel added during the fabrication process [47, 48]. Glucose, oxygen, carbon dioxide, and water are transported between the cells and the supply network. For the model we consider an artificial supply network with only a few assumptions on the specific structure. Mostly, we follow the textbooks [49– 51]. When modeling the tissue and supply network we strike a balance between spatial resolution on one hand and computational effort for the simulation on the other hand. However, the model, or parts of it, could be easily extended to include more detailed descriptions.

We assume that all considered supply networks Ω_*c*_ are composed of connected rigid cylindrical channel segments Ω_*c,i*_ indexed by *i*. We assume fully developed, steady-state and axisymmetric flow of an incompressible medium with low Reynolds numbers and no-slip at the channel wall. The Reynolds number is defined as 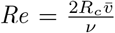, where 2*R*_*c*_ is the inner diameter of the channel, 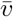 is the mean velocity inside the channel, and ν is the kinematic viscosity of the flowing medium (we omit the index *i* for brevity). In this case, a complex network of channels can be modeled as a graph, where each channel segment is represented by an edge and channels are connected via nodes. At every junction in the network, the volumetric flow rates *Q*_*i*_ as well as the solute flow rates inside the channel segments *i* are given by Kirchhoff’s current law conserving the total flow through the channels following

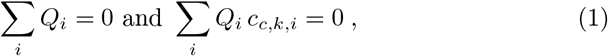

where the summations extend over the channels that are connected to the corresponding junction and *c*_*c,k,i*_ is the concentration of species *k*. Here, the convention is that flow rates into (out of) a channel are positive (negative). In this network, the pressure loss Δ*p*_*i*_ of each channel segment *i* is given by Hagen-Poiseuille’s law

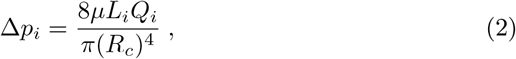

where *µ* is the dynamic viscosity and *L*_*i*_ is the length of the channel segment *i*. This expression relating the pressure drop to the volumetric flow rate is valid for fully developed flow, which translates to low enough Reynolds numbers. Furthermore, we specify the concentrations of the metabolic species *k* at the inlet to the channel network as well as the pressure drop across the network or the flow rate at the inlet.

In the following we usually discuss only one channel segment and therefore often omit the index *i* for brevity. In addition to the assumptions for the flow inside the channels above, we assume that diffusion in axial direction is negligible, which means that convective transport dominates. This is justified for large axial Péclet numbers 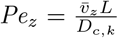. Integrating the resulting convection-diffusion equation over the channel cross sectional area results in an axial transport equation for the mean concentration 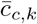 of the relevant species O_2_, CO_2_, and glucose inside each channel segment along the coordinate *z* ∈ [0, *L*]:

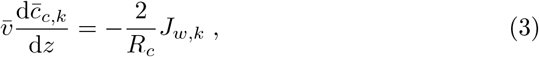

where 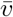 is the average velocity and *J*_*w,k*_ is the diffusive flux at the interface between channel interior and wall given by

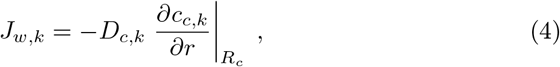

where *D*_*c,k*_ is the diffusion coefficient inside the channel. Further, for small values of the Graetz number 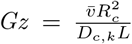, which is the radial diffusion time scale relative to the axial convection time scale, or low species flux through the channel wall (due to slow metabolic reactions or low diffusivity inside the cell-containing tissue compartment), we neglect the radial gradients of species concentration 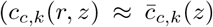 everywhere inside the channel). However, modeling the radial concentration gradients based on mass transfer coefficients would be possible, too.

The concentration in the wall segment Ω_*w*_ is modeled for *R*_*c*_ < *r* < *R*_*c*_ + *t*_*w*_, where *t*_*w*_ is the wall thickness, by

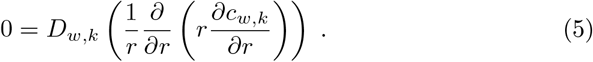

To model the effective diffusion in the porous channel wall with porosity *f*_*p*_ several approaches can be utilized. Here, the effective diffusion coefficients *D*_*w,k*_ are approximated by a simple linear superposition

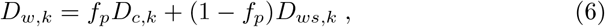

where *D*_*ws,k*_ is the (usually small) diffusion coefficient in the solid part of the channel wall. Such a model would be suitable for a wall material with parallel disconnected pores. However, it would be easily possible to plug-in more complicated models for the effective diffusion coefficients to include tortuosity or pore size effects.

The interface condition between the channel wall and the tissue at the outer channel radius *R*_*w*_ = *R*_*c*_ + *t*_*w*_ is given by

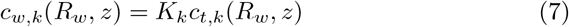

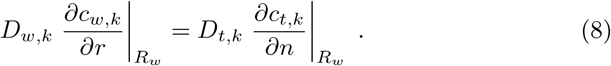

The partition coefficient *K*_*k*_ takes into account that the solutes may have different solubilities in the medium filling the pores and in the cell-containing tissue compartment. It is set to unity in the following, because we assume mainly water inside the channel, pores, and cell-containing tissue compartment, resulting in equal solubilities. However, in situations where a significant solubility contrast is present, we could easily plug-in suitable expressions for the partition coefficients

Finally, the concentration fields in the tissue Ω_*t*_ are modeled by a steady-state reaction-diffusion equation:

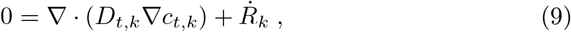

where *D*_*t,k*_ is the diffusion coefficient inside the tissue and 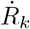 incorporates the metabolism of the cells (reaction rate). We assume that the cellular environment is at rest and therefore the metabolic species are transported in the cell-containing tissue compartment via diffusion only.

The net forward reaction equation for the cellular aerobic metabolism of a well-vascularized tissue is

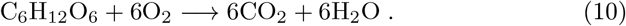

If we assume that the supply of nutrients (such as glucose) to the cells is in abundance compared to that of oxygen, the overall reaction rate can be approximated based on the rate of consumption of oxygen following the Michaelis-Menten kinetics given by

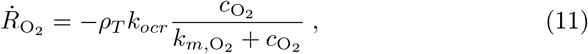

where *ρ*_*T*_ is the number density of cells, *k*_*ocr*_ is the maximum rate of oxygen consumption of the cells in an abundance of oxygen, and 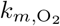 is the Michaelis constant. Note that in addition to the species concentrations, the cell density *ρ*_*T*_ can vary spatially. The reaction rates of the other involved metabolic species apart from oxygen are given in terms of their stoichiometric coefficients from Equation 10.

To further reduce the number of design parameters, we introduce the species flux through the channel wall. This can be calculated from Equation 5 as

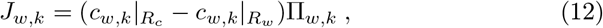

where the permeability of the channel walls is defined by

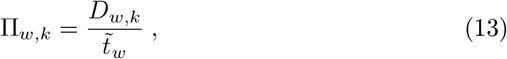

with 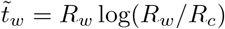 representing the logarithmic wall thickness. The wall permeability Π_*w,k*_ combines the influence of the geometric properties, namely wall thickness, porosity and outer channel radius, with the influence of diffusivity in one single parameter. A variation of Π_*w,k*_ can be achieved by either increasing the diffusivity *D*_*w,k*_ inside the wall material (by changing the material or the porosity) or by decreasing the outer radius or the wall thickness. The permeability of channel wall materials used in several 3D-printing approaches including 2-photon polymerization is typically not high enough to support tissues [45, 52]. To overcome this problem, pores can be 3D-printed in the channel walls covering a fraction *f*_*p*_ of their outer surfaces. On the other hand, in case of supply networks simply consisting of hollow channels resulting from the removal of sacrificial materials [32, 53], we consider an infinite channel wall permeability.

## 3. Design examples

To demonstrate the applicability and possibilities of computational design of artificial supply networks, we simulate several different structures under different conditions. The overall goal is to maintain large human tissue constructs of the order of multiple centimeters over multiple days. Here, we seek supply network designs that fill the available volume, supply the tissue as homogeneously as possible, provide mechanical stability and can be scaled-up. Furthermore, as microfluidic systems are prone to blockage by gas bubbles, clogging, or fouling, the supply network should provide some robustness and redundancy.

Usually, for the design of supply networks it is sufficient to consider the oxygen distribution inside the network and in the cell-containing tissue compartments, while glucose and CO_2_ are neglected. Therefore, in the following we omit the species index *k*. Glucose has a very high solubility in the aqueous culture medium flowing in the supply channels as well as a low cellular metabolic limit concentration. In the case of the reaction product CO_2_, while oxygen is consumed, the same amount of CO_2_ is produced and diffuses through the cell-containing tissue compartment to the network structure. The metabolic limit above which hypercapnia happens is very high, while the diffusive transport through the cell-containing tissue compartment is comparable to that of oxygen. Moreover, the solubility of CO_2_ in aqueous culture media is much higher than that of oxygen, see Table 3. Thus, we can conclude that glucose and CO_2_ do not represent limiting factors in terms of the network design.

Mathematically, we aim to maximize the volume of the cell-containing tissue compartment Ω_*t*_ supplied with oxygen above a concentration threshold (hypoxia) while minimizing the required volume for the network Ω_*c*_ + Ω_*w*_,

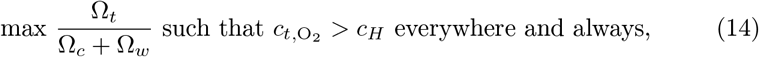

where *c*_*H*_ is the oxygen concentration of hypoxic conditions, see Table 2. Note that other criteria could be easily applied, such as maximizing the mean distance between channels, a maximum concentration of oxygen, or a low spatial variation of concentration.

**Table 1:**
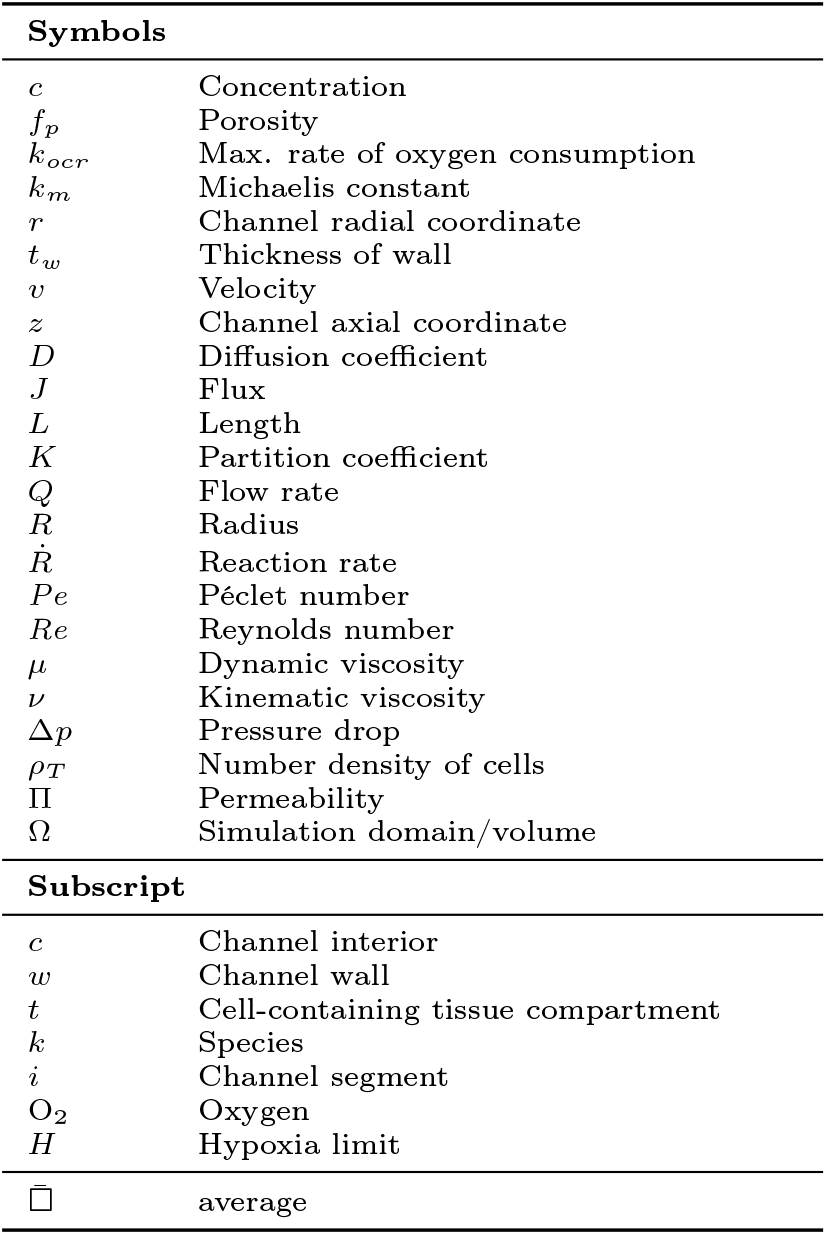
Mathematical notation used in this article.

**Table 2:** Typical data on vascular networks and metabolic activity of various human tissues. The majority is *in vivo* data.

| Parameter | Tissue types |  |
| --- | --- | --- |
|  | High metabolic rate and capillary density | Low metabolic rate and capillary density |
| Type of relevant tissues | Heart, brain, kidney, liver | Fat, bone, cartilage |
| Tissue cell density $\rho_T$ (cells mm<sup>-3</sup>) | $(0.57 \text{ to } 3.3) \times 10^5$ [57–60] | $1.4 \times 10^4$ [61] |
| Maximum oxygen consumption rate $k_{ocr}$ (mol s<sup>-1</sup> cell<sup>-1</sup>) | $1 \times 10^{-16}$ [62, 63] | $(1.2 \text{ to } 4.2) \times 10^{-17}$ [64, 65] |
| Michaelis constant $k_{m,O_2}$ (mol m<sup>-3</sup>) | 0.015 to 0.028 [26, 66] | 0.003 to 0.063 [65, 67, 68] <sup>a</sup> |

<sup>a</sup> Values also include bovine and avian tissue.

**Table 3:** Data for primary species involved in aerobic cellular metabolism.

| Species data | O<sub>2</sub> | CO<sub>2</sub> | Glucose |
| --- | --- | --- | --- |
| Diffusivity in mammalian tissue, $D_t$ ($1 \times 10^{-3} \text{ mm}^2 \text{ s}^{-1}$) | 1.5 to 2.5 [69, 70] | 0.62 to 1.9 [71, 72] | 0.091 to 0.26 [73, 74] |
| Reference concentration in aqueous cell culture medium, $c_{in}$ (mM) | 0.2 to 0.22 [63] | 1.5 to 1.6 [75] | 5.5 to 55<sup>a</sup> |
| Typical cellular metabolic limit, $c_H$ (mM) | < 0.05 (hypoxia) [76, 77] | > 2.4 to 6.5 (hypercapnia) [75, 78] | < 0.03 [79] |

<sup>a</sup> Typical values for commercial mammalian cell culture media.

Exemplarily, we consider different network topologies with realistic oxygen supply scenarios. If not stated otherwise, as an example we assume human liver cells, i.e., high metabolic rate cells, supplied by a typical aqueous cell culture medium. Following the design process depicted in Figure 1 we start by fixing the biological cell parameters, see Table 2 and Table 3: a tissue cell density *ρ*_*T*_ = 2 × 10^5^ cells mm^−3^, oxygen Michaelis constant 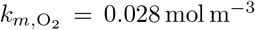, maximum oxygen consumption rate *k*_*ocr*_ = 1.00 × 10^−16^ mol cell^−1^ s^−1^, and a critical oxygen concentration (hypoxia) of *c*_*H*_ = 0.05 mol m^−3^. In addition, Table 2 lists a range of biological parameters for different cell/tissue types which can be applied in such a design process. Next, we consider a typical cell environment with an oxygen diffusion coefficient in the cell-containing tissue compartment of 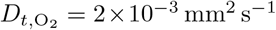. If not stated otherwise, we consider channels with an outer radius *R*_*w*_ = 50 µm and a range of wall permeabilities from 0.05 to 0.2 mm s^−1^. The oxygen diffusivity in the aqueous solution filling the channels is given by 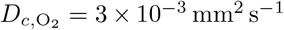. We consider multiple oxygen concentrations at the inlet to the network (0.1 mol m^−3^, 0.2 mol m^−3^ and 1.0 mol m^−3^), which correspond to a rather low oxygen concentration, standard incubator conditions at 37 ^◦^C and 1 atm with 21 Vol% oxygen and 5 Vol% CO_2_, and maximum solubility of oxygen in water at 37 ^◦^C and 1 atm, respectively.

The model from subsection 2.2 is implemented completely open source in *Python* using the finite-element toolbox *FEniCS* [54]; the discretization is performed via the Python-interface of *gmsh* [55]. Most of the following examples are solved as optimization problems approximated via the package *gradient-free-optimizers* [56]. The code implementing the examples is freely available^1^.

### 3.1. Single, spherical cell-containing tissue compartment

As a reference case, we first consider the situation of a single cell-containing tissue compartment with spherical shape, consisting of cells and their environment, submerged into nutrient solutions with different oxygenation levels - from 0.2 mol m^−3^ with 21 Vol% of oxygen at atmospheric pressure, i.e., standard cell culture incubator conditions, to maximal dissolved oxygen of 1.0 mol m^−3^, see Figure 3A. This may appear as a promising configuration due to its simplicity. However, even under ideal conditions (e.g., maximum oxygen concentration and negligible diffusion resistance at the surface of the sphere), the model predicts a maximum possible sphere diameter of around 1.3 mm while keeping the oxygen concentration above the hypoxic cellular limit (Figure 3B). The maximum sphere diameter is greatly reduced for lower oxygen supply concentrations like those in standard incubators, namely 0.2 mol m^−3^. In that case, only a sphere diameter *<* 0.6 mm seems possible (Figure 3B). Similar results, namely a maximal sphere diameter of ≈ 0.7 mm using 0.2 mol m^−3^ oxygen, are obtained with metabolic parameters from [66] (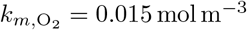, *k*_*ocr*_ = 7 × 10^−17^ mol s^−1^ cell^−1^). These findings underpin the need for a network that allows supplying larger tissues.

**Figure 3:**
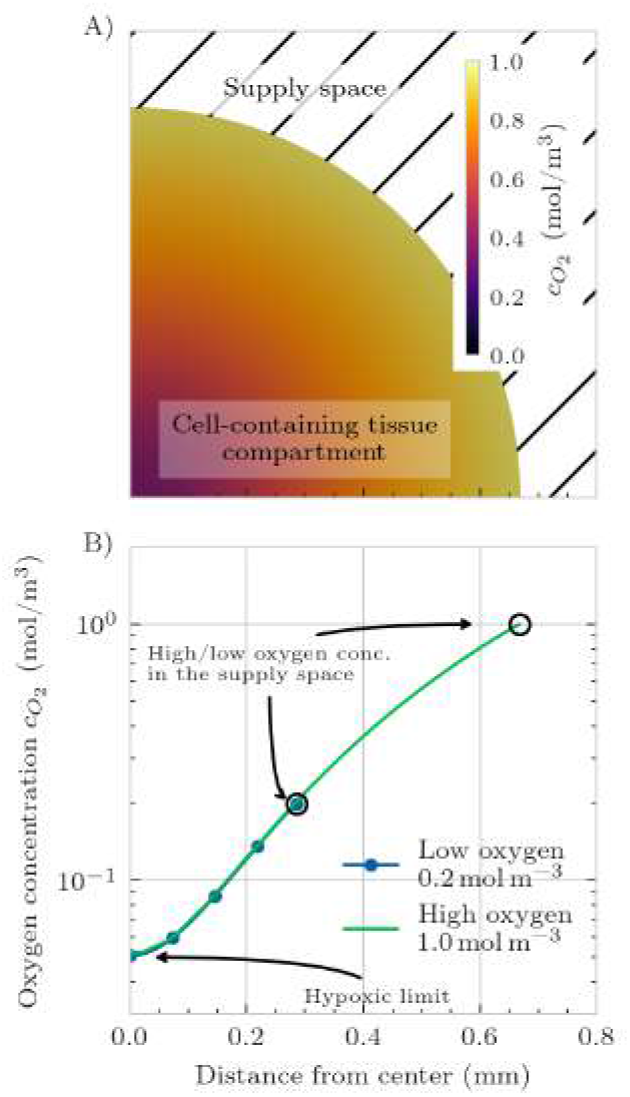
Single spherical cell-containing tissue compartment surrounded by a supply space with an aqueous cell culture medium at constant oxygen concentrations: Oxygen concentration field on a slice through the center of the sphere for the maximum achievable oxygen concentration in the supply space (A) and concentration plot along a line from the center of the sphere for two different concentration levels (B). The hypoxic limit is set to *c*_*H*_ = 0.05 mol m^−3^

### 3.2. Cartesian supply network with cuboid base units

Independent (not interconnected) straight parallel channels running through a tissue provide a scalable and simple alternative to the complex branching and quasi-fractal structures of natural vascular networks [80]. However, this configuration is missing redundancy against blockage and is of low mechanical stability. To provide alternative flow paths in the event of blockages, secondary channels orthogonal to the primary channels can be introduced, which also contribute to the mechanical stability of the network structure and support the cell-containing tissue compartment. Such a supply network can be set up as a Cartesian grid structure with cuboid base units, i.e., a cuboid honeycomb (Figure 4). Each base unit encloses the cell-containing-tissue compartment with the channels located along the edges, see Figure 4C. To supply the channels with oxygen- and nutrient-rich aqueous culture medium, we can place the engineered tissue and network structure between two reservoirs connected to the in- and outlet planes of the structure (Figure 4A). Alternatively, similar to [81], only the lower-right corner can serve as input port, with the outlet port located at the upper-left corner. In all cases, the aqueous solution flows from the inlet reservoir to the outlet reservoir through the artificial supply network. Within the enclosure, a cell-containing tissue compartment (Figure 4C) exchanges metabolic species with the culture medium flowing through the supply network channels. To provide high throughput of oxygen and nutrients between channel and cell-containing tissue compartment, the walls of the supply channels have pores (Figure 4D). Alternatively, using sacrificial templating, channels without dedicated walls can be manufactured [32, 82].

**Figure 4:**
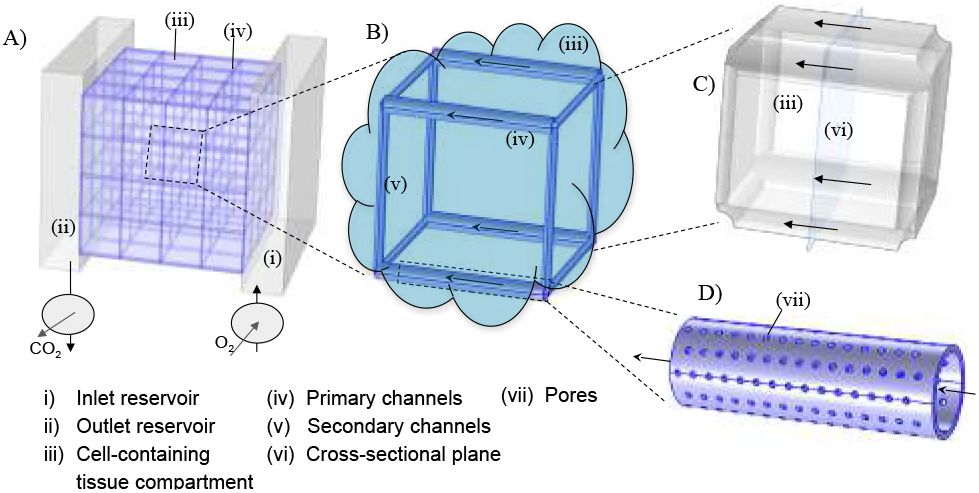
Illustration of a possible configuration of a Cartesian supply network connected to reservoirs through inlet and outlet planes (A). The network is formed by 4×4×4 cuboid base units (B) with channels along the edges enclosing the cell-containing tissue compartment (C). Due to the connection to the reservoirs, the channels in the direction of the main flow (see arrows) serve as primary channels, whereas the channels normal to the flow direction are the secondary channels (here shown with different channel radii). The channels are cylindrical and straight with pores in the walls (D).

To ensure the desired supply of oxygen to the cell-containing tissue compartments even for larger networks, we have to ensure that the velocity inside the defined channel segments is high enough to provide enough oxygen, but low enough to keep the pressure drop within limits. Furthermore, a channel blockage will result in a redistribution of flow through other channel segments. If the velocity is too low, the oxygen concentration in the channels will decrease rapidly over the length of a channel, see Equation 3 and the results below. However, if the velocity is too high, the pressure drop can greatly increase. This can result in high pressure differences between the channel interior and the cell-containing tissue compartment, leading to bypass flow inside the compartment. Using Equation 1 and Equation 2, we can estimate the velocities and pressures inside such networks, see Figure 5. In Figure 5A, simulation results for the network under nominal conditions are presented. The velocity is represented by colored arrows. As expected for the type of inlet/outlet configuration as shown in Figure 4, the flow is driven along the primary channels with a maximum velocity of around 2.5 mm s^−1^. In Figure 5B, we introduced a blockage (dotted black line). In the region around the blockage, flow occurs through secondary channels, too. However, the flow velocity is lower compared to the primary channels. As a consequence, the oxygen concentration inside the channels might get depleted, which would result in lower oxygen concentrations in the cell-containing compartment surrounding the blockage.

**Figure 5:**
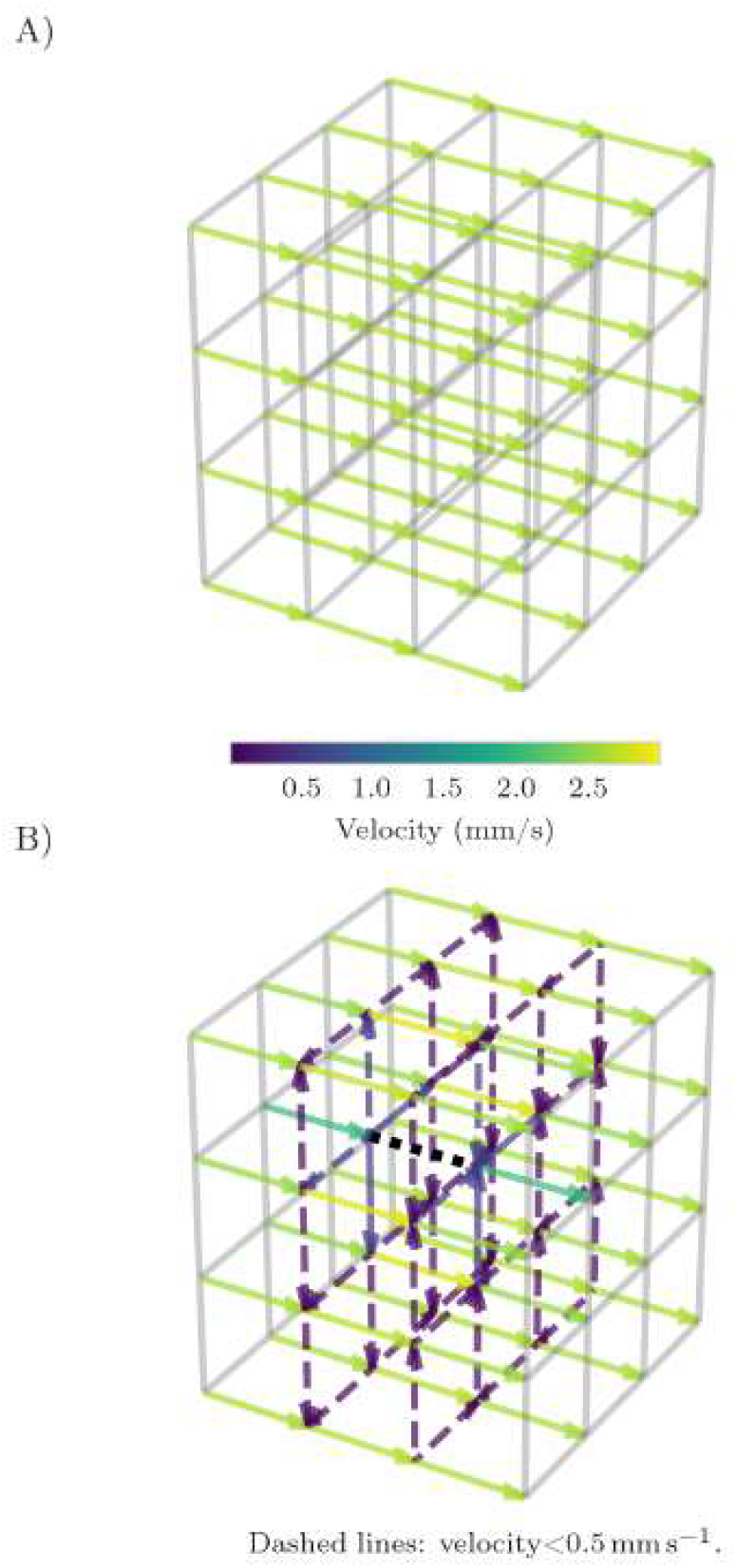
Simulation results for the flow in a supply network under nominal conditions without blockage and flow only through primary channels (A) and with blockage of a single primary channel segment at the center of the network (see dotted black line), leading to a redistribution of the flow through secondary channels (B). The arrows indicate the flow direction; color encodes the flow velocity.

To assess the performance of cuboid supply networks and examine the influence of several design parameters, we present in Figure 6 a selection of simulation results using the model described in subsection 2.2. Here, we impose a fixed oxygen concentration along the channel wall and therefore do not include axial depletion along the flow direction. Axial depletion and its consequences for downstream compartments are addressed separately in Figure 8. At first, we simulate a single cubic base unit with symmetry boundary conditions along its surfaces (Figure 6A). By repeating the base unit in all spatial directions via the symmetry condition the single unit becomes part of a cubic honeycomb . The edge of a cube is 0.5 mm long and we neglect the wall thickness. Here we assume identical flow rates through all channels, with a constant concentration along the outer channel wall. The model predicts the distribution of oxygen, which is high near the channels and decreases towards the center of the cube, with the lowest concentration of oxygen, by design, just above hypoxic condition.

**Figure 6:**
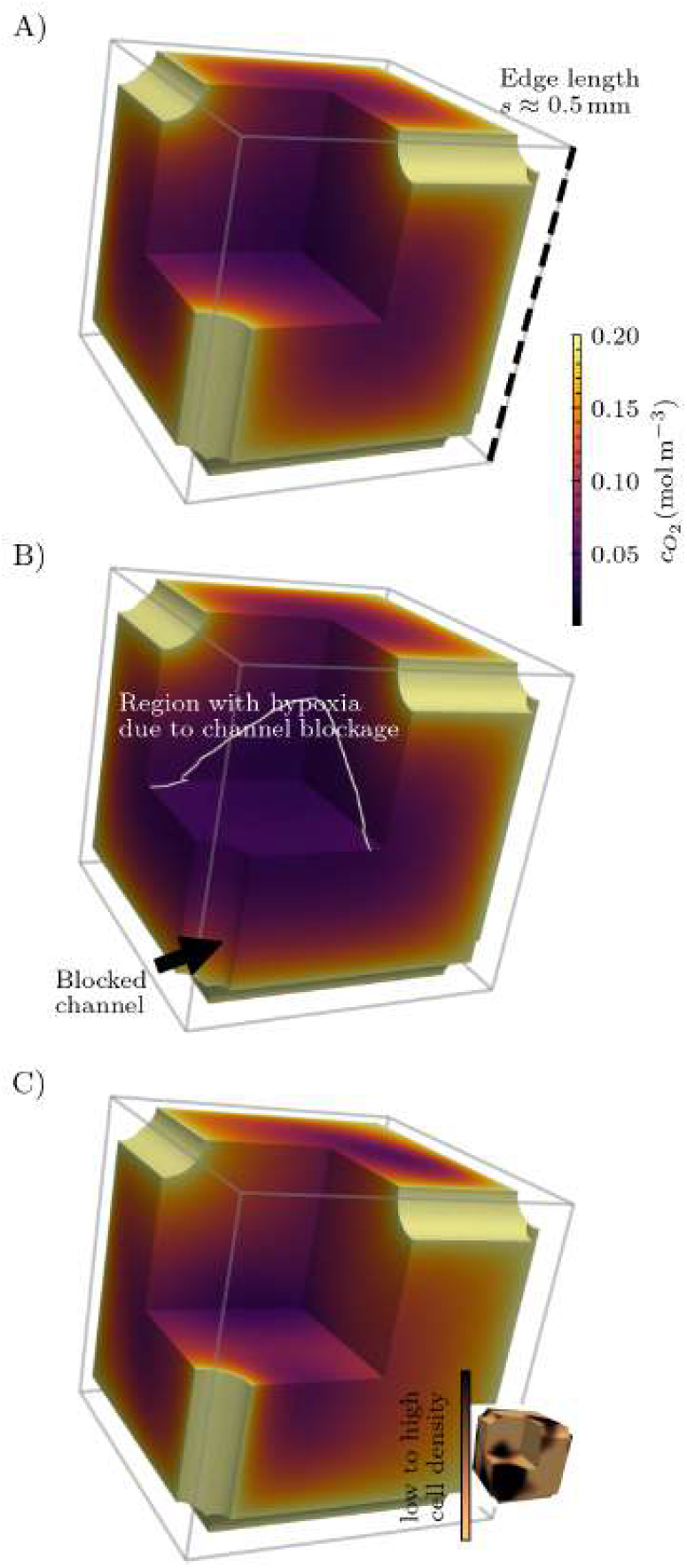
Simulation results of the oxygen distribution inside the cell-containing tissue compartment of a single cuboid base unit with 0.5 mm edge length and symmetry conditions along its surfaces under normal conditions (oxygen concentration outer channel wall 0.2 mol m^−3^, outer channel radius 50 µm); the flow rates through all channels are identical, with a constant concentration along the outer channel wall (wall thickness neglected) (A), with a single blocked channel segment, see arrow, (B), and with a non-homogeneous cell density, see the corresponding cell density distribution in the inset (C).

One intention behind choosing a cuboid design is its robustness with respect to clogging of a channel because of the redundancy of the supply network. Generally, a complete blockage of a channel can lead to critical variations in species supply and, consequently, metabolic activity and cell viability within the cell-containing tissue compartment. Such disturbances may arise, for example, from bubble formation in the channels, a common issue in microfluidic devices operated for extended periods [83]. Therefore, a network should be designed such that the impact of clogging on the metabolic activity does not induce hypoxia in the cell-containing tissue compartment. In Figure 6B the predicted oxygen distribution inside the base unit with a completely blocked channel segment is shown. The encircled area shows the volume inside the cell-containing tissue compartment in which an oxygen concentration below the hypoxic threshold is present. The blocked channel segment is modeled as a surface that is impermeable for oxygen, i.e., the fixed oxygen concentration at the channel wall is replaced by a no-flux condition. In addition, the usually tiny diffusive flux of oxygen through the blocked channel is neglected. Note that for simplicity we simulate only a single base unit with symmetry conditions along its surfaces. Consequently, translated to the entire network, not only one blockage would occur, but many more in all three spatial directions. Owing to the fact that this corresponds to periodically repeating blockages, this is a worst-case scenario when compared to the blockage of only a single channel segment.

Moreover, we can examine the influence of different cell densities on the distribution of oxygen inside the cell-containing tissue compartment. It may be necessary to adapt the operating conditions of the network after an initial cell seeding which is often followed by periods of cell proliferation leading to increased cell densities. While any distribution of cell density, e.g., from experiments, can be considered in the model, here we generate a random distribution of cell density with a spatial correlation obtained from a Gaussian kernel (see the inset in the bottom right of Figure 6C). Note that while in reality the diffusivity inside the cell-containing tissue compartment depends on the cell density distribution, we keep it constant here. It is clearly visible that this inhomogeneous cell density skews the oxygen distribution inside the cell-containing tissue compartment (Figure 6C). Note that in the immediate vicinity of the channel walls, where the highest oxygen concentrations are present, a moderate increase in cell density does not result in a significant reduction in oxygen concentration (Figure 6C).

To identify key parameters for the oxygen supply, we explored the effects of varying geometry parameters of a cuboid network while maximising the cube edge length *s* and preventing oxygen concentrations below the hypoxic threshold. Figure 7 shows the resulting ratio of the tissue compartment volume to the supply network volume, Ω_*t*_*/*(Ω_*c*_ + Ω_*w*_), as a function of the oxygen concentration at the inner channel wall 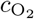, for four combinations of cellular metabolic rate and cell density (panel A: high metabolic rate tissue with high cell density, cellular parameters as applied in all previous simulations, B: high metabolic rate tissue with 1/10 of the cell density applied in A, C: high metabolic rate tissue with high cell density as in A, but lower 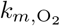, D: low metabolic rate tissue with low cell density), three outer channel radii *R*_*w*_ (0.05 mm, 0.10 mm, and 0.20 mm), and four wall permeabilities Π_*w*_ (0.05 mm s^−1^, 0.1 mm s^−1^, and 0.2 mm s^−1^, plus the limiting “no wall” case representing channels generated by removing sacrificial material). The effect of the wall permeability on the diffusion of oxygen through the channel wall is considered, while the oxygen concentration along the inner channel wall is kept constant. Missing data points at low 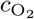 or large *R*_*w*_ correspond to parameter combinations for which no solution satisfying the hypoxia constraint *c*_*H*_ = 0.05 mol m^−3^ everywhere in the tissue compartment could be found. The results in Figure 7 demonstrate that supply networks with different parameter combinations can support cell-containing tissue compartments of similar volume, illustrating how our approach can be used to design a tailored base unit or network structure. By specifying the oxygen concentration inside the channels, which can be adjusted through different laboratory conditions and setups as described in subsection 2.2, the required geometric properties of the network under these conditions are obtained via simulation. Across all four panels, the oxygen concentration at the inner channel wall has the strongest influence on the achievable tissue compartment to network volume ratio. For channels of 50 µm outer radius in panel A, this ratio increases approximately linearly with 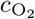, rising from ∼3 at atmospheric oxygen concentration (0.2 mol m^−3^, standard incubator conditions) to ∼10 at 0.6 mol m^−3^ and to more than 13 at the maximum dissolved oxygen concentration of 1 mol m^−3^. Comparable relative gains are observed in panels B, C, and D, although the absolute ratios differ by more than an order of magnitude between high (Figure 7 A–C) and low (Figure 7 D) metabolic rate conditions. The outer channel radius *R*_*w*_ has a markedly weaker influence on the volume ratio than 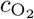. Channels of 100 µm and 200 µm radius consistently yield lower ratios than 50 µm channels under otherwise identical conditions, and the 200 µm curves remain low across most of the simulated range (Figure 7 A–D). Increasing *R*_*w*_ therefore does not translate to a larger tissue compartment per unit network volume; instead, the additional volume associated with wider channels outweigh the modest gain in the cell-containing tissue compartment that can be supplied with oxygen around each channel. The wall permeability Π_*w*_ only has a minor effect on the volume ratio across the simulated range (Figure 7 A–D). For given *R*_*w*_ and 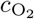, the curves for Π_*w*_ = 0.05 mm s^−1^, 0.1 mm s^−1^, and 0.2 mm s^−1^, together with the limiting “no wall” case, cluster closely together for all conditions (Figure 7 A–D), with differences typically smaller than those introduced by a small change in 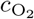 or *R*_*w*_. This indicates that, under the applied parameters, transport across the channel wall is not the rate-limiting step; oxygen supply is instead dominated by diffusion and consumption within the tissue compartment. The absolute volume ratio depends strongly on tissue metabolic demand. For high metabolic rate tissue and high cell density (*ρ*_*T*_ = 2 × 10^5^ cells mm^−3^, 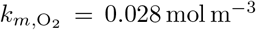 Figure 7A), the ratio of tissue compartment to network volume at atmospheric oxygen concentration and 50 µm radius is approximately 3–4. A ten-fold reduction in cell density (panel B) increases this ratio to approximately 10–15 under the same conditions. Lowering the Michaelis constant 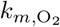 to 0.015 mol m^−3^ while keeping the cell density high (panel C) produces ratios similar to panel A, while a low metabolic rate and low cell density tissue (Figure 7D) (*ρ*_*T*_ = 1.4 × 10^4^ cells mm^−3^, 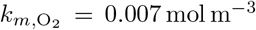, *k*_*ocr*_ = 4.2 × 10^−17^ mol s^−1^ cell^−1^) yields the highest ratios overall, exceeding 300 at 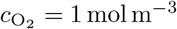 and *R*_*w*_ = 0.05 mm.

**Figure 7:**
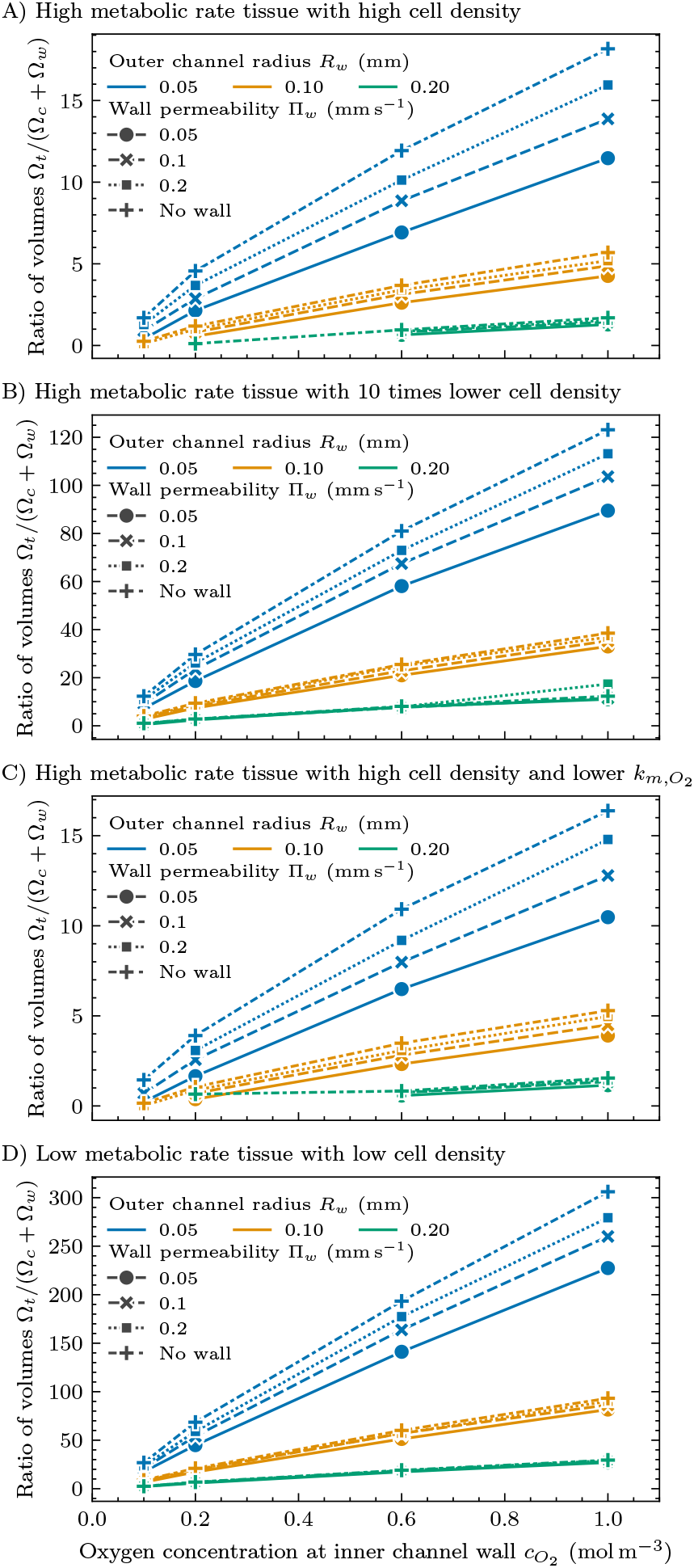
Simulation results of the influence of design parameters including wall permeability and oxygen concentration at the inner channel wall on the maximum base unit size that keeps the oxygen concentration everywhere inside the cell-containing tissue compartment above the hypoxic limit of *c*_*H*_ = 0.05 mol m^−3^ for different metabolic conditions (A: *ρ*_*T*_ = 2 *×* 10^5^ cells mm^−3^, 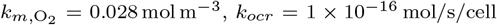, *k*_*ocr*_ = 1 *×* 10^−16^ mol*/*s*/*cell; B: identical to A but with *ρ*_*T*_ = 2 *×* 10^4^ cells mm^−3^; C: identical to A but with 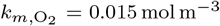; D: *ρ*_*T*_ = 1.4*×*10^4^ cells mm^−3^, 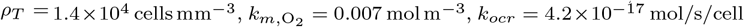, *k*_*ocr*_ = 4.2*×*10^−17^ mol*/*s*/*cell). “No wall” refers to a nearly infinite channel wall permeability, representing channels generated by removing sacrificial material. The lines are guides to the eye. Missing data points indicate no valid solution.

**Figure 8:**
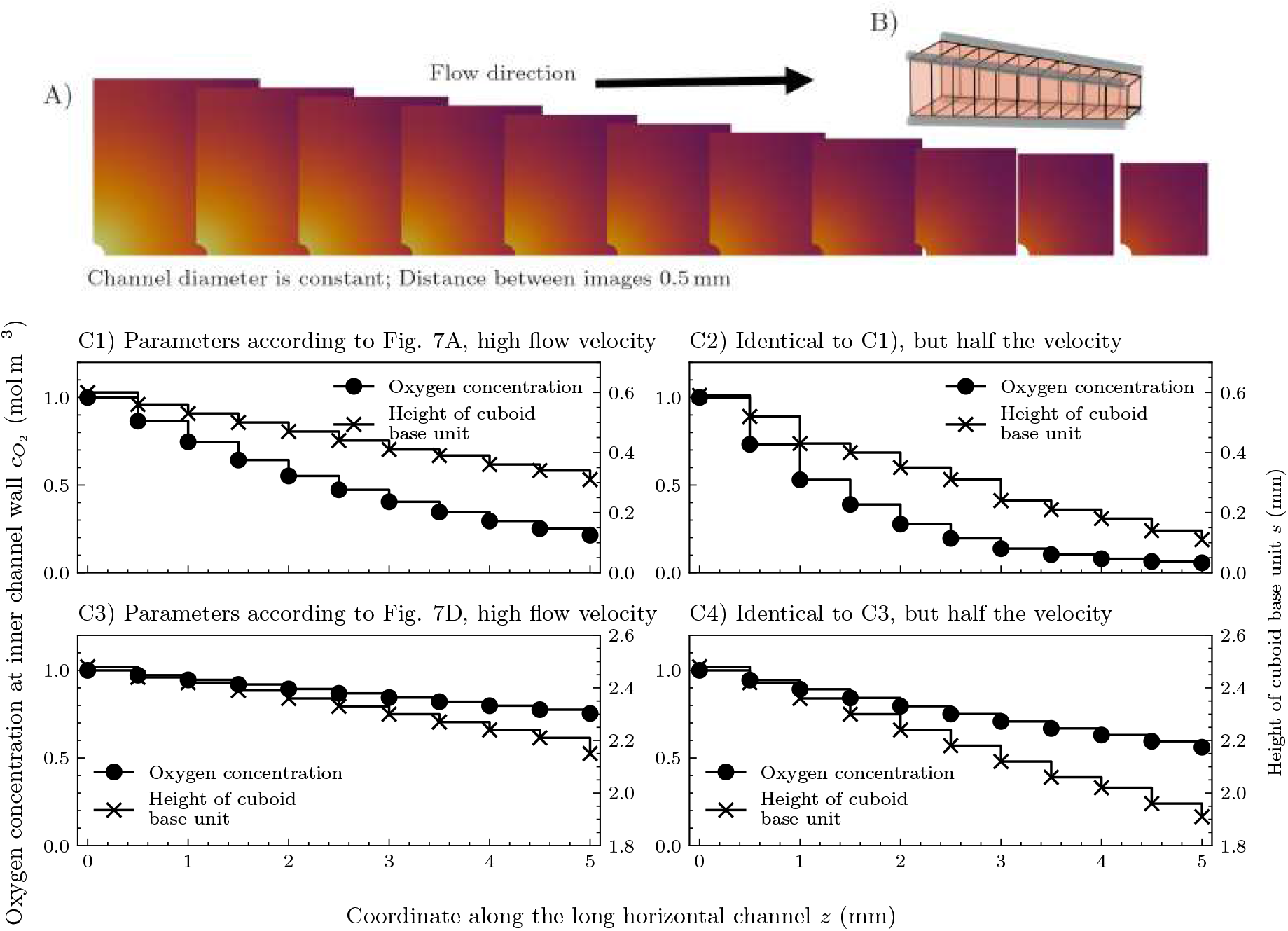
Simulation result of the oxygen concentration inside a long channel with different flow velocities and the resulting reduction of maximum possible cuboid base unit size along the channel length to prevent hypoxia (*c*_*H*_ = 0.05): Visualization of the oxygen distribution inside the cell-containing tissue compartment at different axial coordinates along the channel with the parameters identical to Figure 7A and a velocity of 8 mm s^−1^ (A), an illustration of the resulting converging channels (B), and the corresponding concentration inside the channel (C1, dots) and the resulting edge length of the cuboid base units to prevent hypoxia (C1, crosses). Similarly, C2–C4 display concentration and resulting edge length along the channel length for different conditions (C2 identical to C1 but half the velocity, C3 identical to Figure 7D and velocity identical to A, C4 identical to C3 but half the velocity).

In case of long network channels, the oxygen concentration of the aqueous cell culture medium inside the channels might get reduced significantly along the length of the channel. Furthermore, as evident from Figure 5, a channel blockage and subsequent rerouting of flow through secondary channels might greatly reduce the flow velocity leading to higher oxygen depletion, too. To examine this effect on the largest possible cuboid base unit size while still preventing hypoxia, we have simulated a long channel segment of 5 mm length without any channels perpendicular to the long channels, see the illustration in Figure 8C. We utilized our model from subsection 2.2 including Equation 3 and the same cell parameters as initially applied and as also used in simulations shown in Figure 7A. Due to the symmetry conditions of the domain, see the illustration Figure 8C, only a quarter of the channel and cell-containing tissue compartment is simulated. The flow velocity inside the long channel is set to 8 mm s^−1^, which corresponds roughly to a Reynolds number *Re* ≈ 1, and the inlet oxygen concentration is the maximum possible concentration of 1.0 mol m^−3^ at 37 ^◦^C.

The channel length is discretized into 10 elements, with the flow direction inside the channel from left to right (see Figure 8A). Using the inlet concentration as the boundary condition at the inner channel wall of the first element we calculate the concentration distribution inside the first element (and maximize the height of the cell-containing tissue compartment while still preventing hypoxic conditions). Then we calculate the flux through the boundary between channel and cell-containing tissue compartment via Equation 4 and solve the discretized Equation 3 for the concentration at the boundary between the channel and the next element. This is repeated step by step until the end of the channel resulting in the step-like plots of inner channel wall concentration and height in Figure 8B). Note that this discretization and solution process introduces small numerical errors, which can be reduced even further by introducing more discretization elements.

Because oxygen diffuses from the channel into the cell-containing tissue compartment, where it is consumed, the wall concentration drops monotonically from inlet to outlet (Figure 8). To prevent hypoxia everywhere, the maximum allowable edge length of the cell-containing tissue compartment normal to the channel must therefore be continuously reduced along the flow direction, producing a tapering compartment (Figure 8B). Figure 8 C1 to C4 quantify this tapering for four combinations of metabolic activity and flow velocity. Figure 8 C1 and C2 correspond to the high metabolic rate, high cell density tissue of Figure 7A, and C3 and C4 to the low metabolic rate, low cell density tissue of Figure 7D. Within each pair, the first panel uses a velocity inside the channel of *v* = 8 mm s^−1^ and the second uses *v* = 4 mm s^−1^. In C1 the concentration in the channel falls from 1.0 to around 0.2 mol m^−3^ over 5 mm, and the permissible edge length shrinks from 0.6 mm to 0.3 mm. In C3 the same channel delivers oxygen to a much less demanding tissue: the wall concentration drops only to 0.7 mol*/*m^3^, and the edge length tapers only from 2.4 to 2.2 mm. Low metabolic rate tissues therefore tolerate much longer channels before axial depletion forces significant tapering. In the high activity case (C2: high metabolic rate with lower velocity), the edge length collapses to near zero before the end of the channel, so a 5 mm channel cannot supply any tissue over its full length without inducing hypoxia. In the low activity case (C4), halving the velocity reduces the final edge length from 2.4 to 1.9 mm, which is only a modest effect, because the channel is far from depletion in either case. Velocity is therefore a critical design parameter for high-demand tissue and only a weak lever for low-demand tissue.

### 3.3. Rhombic dodecahedral base unit

One disadvantage of cubic base units is the large ratio of minimal and maximal distance between a point at the center and points along the edges where the channels are located. This results in a relatively high variance in oxygen distribution, see Figure 6A. However, in addition to the cuboid honeycomb geometry, several other volume-filling, scalable designs are possible. Here we present one of these structures as an example: the rhombic dodecahedral honeycomb provides a smaller difference between the minimal and maximal center-to-edge distance, see Figure 9, and can be readily examined using our model and design approach. Similar to Figure 6A we neglect the effect of the wall thickness. Figure 9A displays the resulting oxygen distribution inside a rhombic dodecahedron with 0.4 mm channel segments along the edges. Under identical hypoxic thresholds the possible volume of the cell-containing compartment is around 50% higher compared to a cubic base unit (compare Figure 6A with Figure 9A).

**Figure 9:**
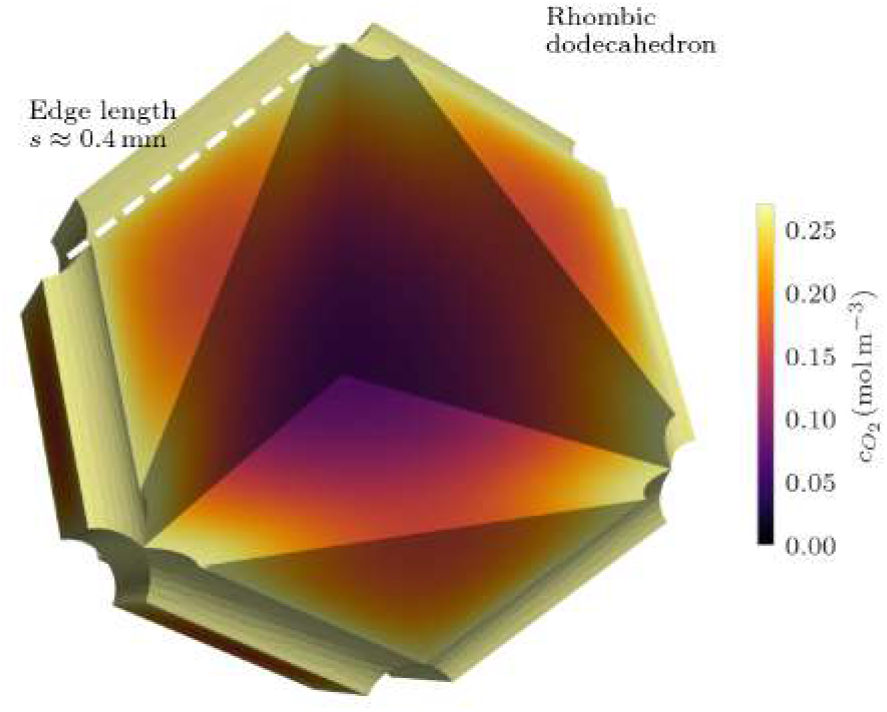
Simulation result predicting the oxygen concentration distribution in a rhombic dodecahedron base unit of a supply network (oxygen concentration at outer channel wall 0.2 mol m^−3^, outer channel radius 50 µm).

## 4. Discussion and Conclusions

The structure of artificial supply networks and the supply of metabolic species strongly influence the cellular metabolic activity and viability of cells within engineered human tissue. However, design strategies to ensure a sufficient oxygen and nutrient supply, as well as waste removal, of constructed human tissue as a whole are largely missing. The novel, systematic, computer-assisted design process presented in this study, using human liver cells as the main example, combines mathematical models of fluid dynamics, cell metabolism, and network properties to identify key parameters influencing the supply performance. Further, such a design process reduces the number of prototypes and experiments needed to identify a suitable or even optimal solution. The mathematical description of the coupled system consisting of cells and their environment in the cell-containing tissue compartment, the channel walls, and the channel interior was implemented in a freely available Python package to facilitate easy utilization and adaption by others. We demonstrated the applicability and possibilities of computationally designing artificial supply networks based on several examples, showing how to easily and quickly explore the vast parameter space consisting of possible network structures, including channel dimensions, channel wall permeability, oxygen concentrations in aqueous cell culture media, network topologies (cuboid and rhombic dodecahedron base units), cell densities, as well as normal and impaired flow conditions.

At first, we examined the most straightforward configuration: a single, spherical cell-containing tissue compartment submerged in an oxygenated nutrient solution. We found that the maximum possible diameter of the sphere while still preventing hypoxia is below 1.3 mm at maximum dissolved oxygen concentrations and below 0.6 mm under standard incubator conditions. The latter results are quite similar to published experimental data on the diameters of human liver cell cultures in spheroid form with and without dead cells in the center, see [84, 85], although the parameters we applied, including cell density, are not specified in these publications or differ slightly. Therefore, a supply space around a spherical cell-containing tissue compartment is not suitable to grow and maintain large human tissues.

As a first realization of a supply network for large tissue constructs, we considered cuboid honeycomb structures. A network model based on the Hagen-Poiseuille velocity field in each channel segment allows computing the flow distribution inside the network. Here, we examined the robustness with respect to the total blockage of a single channel segment that might for example occur upon bubble formation. These examples provide vital information needed to design network structures taking redundancy and blockage into consideration. Nevertheless, certain fabrication and operational precautions should be taken to prevent bubble formation inside the channels. Bubbles usually form at nucleation sites at the channel wall, a process that is influenced by the surface properties of the channels. To reduce the likelihood of bubble formation and growth, care should be taken during fabrication to avoid sharp edges and recesses at the channel walls. In addition, well-wettable wall materials are preferred.

Obtaining the largest possible base unit forming a structured network is an important aim in human tissue engineering. Therefore, we computed the oxygen concentration field in a cubic base unit for several points in the parameter space, while maximizing the size of the cell-containing tissue compartment within the base units. We found that the oxygen concentration inside the channels at the inner channel wall has the largest impact on the achievable tissue compartment to network volume. Further, a low outer channel radius and high wall permeability contributed to high volume ratios between the cell-containing tissue compartment and the supply network, too. If the networks get larger in total size and therefore longer channels are required, the oxygen concentration inside the aqueous culture medium significantly reduces along the channel length. We examined this effect by simulating long channels of 5 mm length using our coupled model. Our results demonstrate that, to prevent hypoxia, the height of the cell-containing tissue compartment normal to the long channel has to be continuously reduced along the flow direction. The severity of this reduction is governed by the interplay of metabolic demand and flow velocity. For high metabolic rate, high cell density tissue, halving the velocity renders a 5 mm channel effectively unusable, whereas the same velocity reduction barely affects low metabolic rate tissue. Velocity selection is therefore far more consequential for high activity and high density tissues.

In addition to the cuboid honeycomb geometry, several other space-filling designs with channels arranged along the edges of base units are possible. As an example, we examined rhombic dodecahedrons. We found that under identical conditions with the same minimal oxygen concentration allowed at any position within the cell-containing tissue compartment, the total volume of the dodecahedron base unit is around 50 % higher compared to a cuboid base unit. Our model also allows incorporating inhomogeneities, especially inhomogeneous cell densities that may occur during the culturing of human tissue.

Our results highlight the importance of considering the structure of the artificial supply network, the effects of non-uniform cell density, channel blockage, and long channels on the oxygen distribution inside the cell-containing tissue compartment. Synthesizing the findings across these examples, the key results and design guidelines are:

- **The oxygen concentration at the channel wall is the strongest design lever**. Across all simulated conditions, increasing 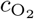 produced approximately linear gains in the achievable tissue compartment-to-network volume ratio for high metabolic rate tissue, and superlinear gains for lower-demand tissue. Increasing 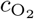 was the single most effective lever in all cases.
- **Smaller outer channel radii supply more tissue per unit network volume**. Channels of 50 µm radius consistently outperform 100 and 200 µm channels in maintaining a high tissue compartment volume. Although larger channels may be easier to fabricate, their volume increases more rapidly than the tissue section that can be oxygenated around them.
- **Wall permeability is not the rate-limiting step**. For all simulated values of permeability (smallest simulated value 0.05 mm s^−1^) including the limiting “no wall” case, permeability only has a minor effect on the volume ratio; the bottleneck is diffusion within the tissue compartment itself. High-permeability walls or even sacrificial templating are therefore secondary to network geometry and inlet oxygenation.
- **Velocity sensitivity is closely related to metabolic demand**. For high metabolic rate tissue, the tolerable channel length is highly sensitive to flow velocity. Halving the velocity can turn a usable long channel into one that gets fully depleted from oxygen before its end.
- **Design for redundancy**. Secondary channels orthogonal to the primary flow direction mitigate the impact of individual blockages.
- **Design for oxygen depletion**. Long channels may exhibit significant oxygen depletion, which can be counteracted by providing a tapered network geometry along the flow direction.
- **Matching the design to the tissue’s metabolic demand**. For high metabolic rate tissue, the network occupies roughly a quarter of the total volume at atmospheric oxygenation. Low-metabolic-rate tissue requires only a fraction of that.

Utilizing our numerical simulations of coupled human tissue models offers a fast and easy way to explore the design space of supply networks. The design process can greatly reduce the vast parameter space to be explored in experiments and provide promising proposals for large supply structures tailored to maintaining specific engineered human tissue under different culture conditions and cell states. This approach has the potential to significantly contribute to the development of more efficient and effective tissue engineering strategies, enabling the creation of large-scale, three-dimensional human tissues that can be used for various applications in therapeutic tissue engineering, disease modeling, and drug testing.

However, the predictions of our model are only as accurate as the assumptions and approximations underlying it. In particular, during the derivation of the model we made several simplifying assumptions concerning physics and biology. Therefore, a minimal number of prototyping and experiments is still required to assess the validity of the model, and experimental validation of the designed networks will be essential to confirm the validity of the design process. To improve the model, further refinements are possible such as diffusion coefficients that depend on the cell number density. Future studies should focus on further refining the design process, incorporating additional biological and physical parameters, and exploring the potential of this approach for other tissue types and applications. Overall, this study demonstrates the power of computational design in tissue engineering. We deem the computational design of artificial supply networks for large human tissue constructs still at the beginning of an impactful development, with the potential to advance the field of tissue engineering and regenerative medicine.

## CRediT authorship contribution statement

**Henning Bonart:** Investigation; Methodology; Software; Visualization; Writing - original draft; Writing – review & editing. **Pramodt Srinivasula:** Investigation; Methodology; Visualization; Writing - original draft. **Ulrike A. Nuber:** Conceptualization; Funding acquisition; Investigation; Supervision; Writing – original draft; Writing – review & editing. **Steffen Hardt:** Conceptualization; Funding acquisition; Methodology; Project administration; Resources; Supervision; Validation; Writing - original draft; Writing - review & editing.

## Data Availability Statement

Data and code is available from the authors upon reasonable request.

## Footnotes

1 The newly developed code that was used to perform the simulations as well as generate the figures is available for the reviewers upon request and will be made available to the public after our manuscript is accepted for publication.

